# Music-selective cortex is sensitive to structure in both pitch and time

**DOI:** 10.1101/2021.12.17.473232

**Authors:** Dana Boebinger, Josh H. McDermott, Nancy Kanwisher, Sam Norman-Haignere

**Affiliations:** Speech and Hearing Bioscience and Technology, Harvard University, Cambridge, MA 02138; Brain and Cognitive Sciences, Massachusetts Institute of Technology, Cambridge, MA 02139; McGovern Institute for Brain Research, Massachusetts Institute of Technology, Cambridge, MA 02139; Department of Biostatistics & Computational Biology, University of Rochester, Rochester, NY 14642; Department of Neuroscience, University of Rochester, Rochester, NY 14642; Center for Brains, Minds, and Machines, Massachusetts Institute of Technology, Cambridge, MA 02139; Zuckerman Institute for Brain Research, Columbia University, New York, NY 10027

**Keywords:** auditory cortex, fMRI, music, melody, rhythm

## Abstract

Music is distinguished from other natural sounds by the presence of relatively discrete notes, which are then organized across pitch and time to convey melody, harmony, and rhythm. Growing evidence suggests that small clusters of neural populations within anterior and posterior human non-primary auditory cortex respond selectively to musical structure. However, it is unclear whether this selectivity reflects short-term musical structure at the level of individual notes, and/or the patterning of notes in pitch and time. We used fMRI voxel decomposition to measure the response of music-selective and non-selective auditory neural populations to synthetic music and drum stimuli whose notes were scrambled in pitch and/or time, disrupting musical pattern structure while largely preserving note-level structure. We observed reliably stronger responses to music with intact pitch and temporal pattern structure in both anterior and posterior music-selective regions bilaterally, but little difference between intact and scrambled music in non-selective populations. Further, only music-selective populations showed reliably stronger responses to note-scrambled music compared with non-music sounds. These results suggest that musical structure involving both individual notes and their patterning over time is specifically represented in localized music-selective neural populations of human non-primary auditory cortex.

## INTRODUCTION

Music is a quintessentially human artifact with distinctive acoustic and structural properties. One defining feature is the presence of discrete “notes” characterized by relatively stable pitch contours and spectral timbres, as well as characteristic temporal envelopes (e.g., attack, decay, sustain, release) (Agus et al., 2012; Lindblom & Sundberg, 2007; McAdams et al., 1995; Ogg et al., 2017; Ozaki et al., 2024; Savage et al., 2015). These acoustic properties substantially affect listeners’ perception of musicality. For example, speech syllables with more sustained pitch contours are perceived as more song-like (Sankaran et al., 2024), especially when repeated (Tierney et al., 2013, 2018).

Beyond individual notes, music is also defined by how notes are structured and organized across pitch and time. We use the term “note-pattern structure” to refer to these relationships among notes that convey melody, harmony, rhythm, and tonality. Tonality, for instance, is conveyed by statistical structure computed across many notes, including both the set of pitches used as well as the statistical regularities with which those pitches occur and co-occur over time (Temperley, 2008). Such patterning is highly structured across musical cultures (Mehr et al., 2019; Savage et al., 2015), with pitches typically drawn from a small, stereotyped scale (Krumhansl & Kessler, 1982; Mehr, 2024). Temporal patterning is likewise strongly periodic (Mehr, 2024; Mehr et al., 2019; Savage et al., 2015), and often organized into complex metrical hierarchies, which can be conveyed even in the absence of strong pitch information (e.g., drumming). Listeners can implicitly learn these regularities, even without formal musical training (Bigand & Poulin-Charronnat, 2006; Rohrmeier et al., 2011), and often quite early in development (Hannon & Trainor, 2007).

Understanding how the brain encodes musical structure is a central challenge of auditory and cognitive neuroscience (Di Liberto et al., 2020; Koelsch et al., 2005; Leaver & Rauschecker, 2010; Norman-Haignere et al., 2015; Peretz et al., 2015; Sankaran et al., 2024). Animal studies have identified tuning for key acoustic primitives, including pitch (Bendor & Wang, 2005; Bizley et al., 2010), pitch contour (Kilgard & Merzenich, 2002; Yin et al., 2008, 2014), interval timing (Merchant et al., 2013), and spectrotemporal modulation (DeCharms et al., 1998; Depireux et al., 2001; Miller et al., 2002; Woolley et al., 2005). Non-human animals also show enhanced responses to surprising notes (Bianco et al., 2024; Yaron et al., 2012) and can learn artificial grammars (Wilson et al., 2013), which may be precursors to the sensitivity to musical pattern structure observed in the human brain (Di Liberto et al., 2020; Sankaran et al., 2024). Growing evidence suggests that the human brain has additional adaptations for processing music-specific structure (Landemard et al., 2021). In particular, neural populations in anterior and posterior non-primary auditory cortex respond more strongly to music than to other sounds (Angulo-Perkins et al., 2014; Boebinger et al., 2021; Leaver & Rauschecker, 2010; Norman-Haignere et al., 2015; Norman-Haignere, Feather, et al., 2022), including synthetic stimuli matched to music in spectrotemporal modulation statistics (Norman-Haignere & McDermott, 2018). Initial evidence came from a statistical decomposition of fMRI responses to natural sounds (Norman-Haignere et al., 2015), and has been replicated in additional fMRI cohorts (Boebinger et al., 2021) and confirmed by intracranial recordings (Norman-Haignere, Feather, et al., 2022). Nonetheless, the role of these neural populations in encoding different types of musical structure remains unclear.

A major open question concerns the type of musical structure represented in auditory cortex. Neural populations in non-primary auditory cortex integrate information over roughly 200-500 ms (Norman-Haignere, Long, et al., 2022; Overath et al., 2015), consistent with music selectivity emerging within a few hundred milliseconds after onset (Leaver & Rauschecker, 2010; Norman-Haignere, Feather, et al., 2022; Norman-Haignere, Long, et al., 2022). This suggests sensitivity to relatively short-term musical structure, possibly reflecting selectivity for individual notes. In contrast, studies using encoding models have found evidence for responses to surprising notes in auditory cortex (Di Liberto et al., 2020) that are specific to music (Sankaran et al., 2024), suggesting some sensitivity to pattern structure. Thus, it remains unclear whether music-selective neural populations predominantly encode note-level acoustic structure or the broader patterning of notes in pitch and time.

A second question concerns the functional and anatomical organization of musical note and pattern representations. Anterior and posterior regions of auditory cortex differ in their spectrotemporal modulation tuning (Hamilton et al., 2018; Hullett et al., 2016a; Norman-Haignere et al., 2015; Santoro et al., 2014a; Schönwiesner & Zatorre, 2009) and in sensitivity to pitch (Norman-Haignere et al., 2013, 2015). A striking property of music-selective neural populations is that they show two distinct, widely spaced clusters in anterior and posterior auditory cortex, raising the question of whether these clusters differ in selectivity for temporal vs. pitch structure. Prior work has also proposed that there is hemispheric specialization for different spectrotemporal modulation rates (Albouy et al., 2020; Zatorre & Belin, 2001), with the left hemisphere biased towards fast temporal processing and the right hemisphere biased towards pitch and melody. While music selectivity appears to be largely bilateral, prior work measuring music selectivity has focused on relatively short sounds (e.g., 2 seconds) and has not explicitly manipulated temporal and pitch structure.

Here, we used a scrambling approach to examine neural selectivity, inspired in part by prior work in high-level visual cortex (Liu et al., 2010). Our objective was to determine whether music-selectivity predominantly reflects individual musical notes or the sequential patterning of notes over time. Music selectivity is often characterized as a preferential response to music compared with sounds that lack canonical musical structure. This selectivity could plausibly arise from sensitivity either to individual musical notes, which have distinctive acoustic properties such as relatively flat pitch contours and characteristic temporal envelopes, or to the higher-order patterning of notes in pitch and time that gives music its melodic and rhythmic character. Scrambling notes in a musical score provides a principled test of this question, as it preserves the acoustic structure of individual notes while disrupting higher-order sequential organization. If music selectivity primarily reflects sensitivity to note-pattern structure, responses should be substantially stronger for intact music than for scrambled music. In contrast, if music selectivity primarily reflects sensitivity to the acoustic properties of individual notes, responses to scrambled music should still be reliably stronger than responses to non-music stimuli. These two non-exclusive hypotheses make distinct, testable predictions regarding the pattern of responses across intact music, scrambled music, and non-music stimuli. To test these predictions, we used functional MRI to measure human auditory cortical responses to synthetic MIDI-generated music clips that were scrambled in pitch and/or time, thus removing virtually all pitch and temporal pattern structure while largely preserving note-level structure. We measured responses to MIDI drum stimuli, which lack strong, fine-grained note-level pitch or harmonic structure, and temporally scrambled them to selectively disrupt temporal patterning. We also tested several non-music stimuli to examine whether responses to the note-scrambled stimuli were still greater than those for other types of sounds. Using component analyses developed in prior work (Norman-Haignere et al., 2015), we isolated music-selective neural populations and compared their responses to those of nearby neural populations that respond selectively to other sound categories or acoustic features. We repeated these analyses in both anterior and posterior auditory cortex, as well as in the left and right hemispheres, to investigate any anatomical or functional organization.

## METHODS

### Participants

Fifteen young adults (7 self-identified female, 8 self-identified male, 0 self-identified non-binary; mean age = 25.6 years, SD = 4.5 years) participated in the experiment. Of these participants, 12 had some degree of musical training (mean = 10.1 years of training, SD = 6.7). All participants were fluent English speakers and reported normal hearing. The study was approved by the Committee on the Use of Humans as Experimental Subjects at MIT (COUHES #1507140467), and written informed consent was obtained from all participants. Participants were compensated with an hourly wage for their time.

### Overview of experimental design

For the current study, participants completed multiple scanning sessions, which included a set of fMRI “localizer” scans to estimate each participant’s voxel weights for the components from our previous study (Norman-Haignere et al., 2015), and the main fMRI experiment in which responses were measured to different music conditions. For the component localizer, participants were scanned while they listened to a set of 30 natural sounds. Based on the voxel responses to these sounds, we then estimated each participant’s voxel weights that best approximated the previously inferred response profiles from the original study (Norman-Haignere et al., 2015). We then applied these weights to participants’ voxel responses in the main experiment, in which they listened to synthetic music clips that had been manipulated by scrambling either the note pitches and/or onset times. This process allowed us to examine how the response components from our previous study (Norman-Haignere et al., 2015) respond to the new stimulus conditions, and thus the extent to which neural selectivity for music reflects note-level vs. pattern structure.

### Component localizer

#### Overview of voxel decomposition

Because a single fMRI voxel pools responses from hundreds of thousands of neurons, voxel responses may be modeled as the sum of the responses of multiple underlying neuronal populations. In our previous work, we found that voxel responses to natural sounds in auditory cortex can be approximated by a weighted sum of six canonical response profiles, or “components:”

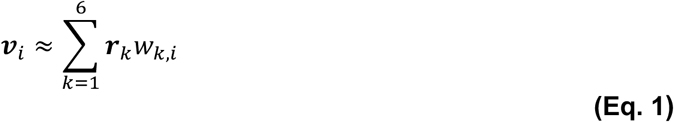

where 𝒗*_i_* is the response (a vector) of a single voxel to the sound set, 𝒓*_k_* represents the 𝑘^th^ component response profile across sounds, which is shared across all voxels, and 𝑤*_k,i_* represents the weight (a scalar) of component 𝑘 in voxel 𝑖. If we concatenate the responses of many voxels into a sound x voxel data matrix, 𝐷, this approximation corresponds to matrix factorization:

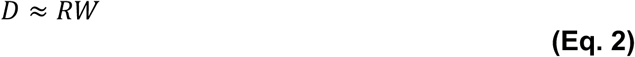

where 𝑅 is a sound x component response matrix and 𝑊 is a component by voxel weight matrix. The component responses and weights were jointly inferred in our prior studies by maximizing the non-Gaussianity of the weights across voxels, akin to standard algorithms for independent components analysis (ICA). The voxel decomposition method is described in detail in our previous paper (Norman-Haignere et al., 2015) and code is available online (https://github.com/snormanhaignere/nonparametric-ica).

Deriving components from scratch requires a large amount of data because both the component responses (𝑅) and weights (𝑊) need to be inferred, which requires using statistics computed across thousands of voxels (requiring multiple participants). However, if the component responses are known (e.g., from a prior study), the voxel weights can be inferred in a single new participant using linear regression (separately for each voxel). In addition, once the component voxel weights are known, the component responses to a new set of sounds, measured in the participants for whom the weights are known, can also be easily inferred using linear regression (separately for each sound). This latter procedure is analogous to identifying a set of voxels that respond to a particular ‘localizer’ contrast, and then measuring their response to a new set of stimuli.

#### Stimuli

To reduce the amount of scan time needed to infer the component weights, we chose a subset of 30 sounds from our original 165-sound set that was best able to identify the six components. These sounds were selected by greedily discarding sounds so as to minimize the expected variance of the inferred component weights. Assuming isotropic voxel noise, the expected variance of the weights, inferred by ordinary least-squares (OLS) regression, is proportional to (Dale, 1999):

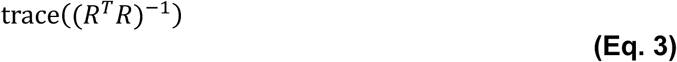

where 𝑅 is the sound x component response matrix. Intuitively, the optimization chooses sounds that have high response variance for each component (e.g., lots of music and non-music sounds for the music-selective component), and where that response variance is relatively uncorrelated across components. Each column/component of 𝑅 was z-scored prior to performing our greedy optimization (which amounts to iteratively discarding rows of 𝑅 to minimize the equation above).

#### Stimulus presentation and scanning procedure

The data for the component localizer came from previous studies on the same participants (n = 2 from Norman-Haignere et al., 2015; n = 7 from Boebinger et al., 2021; and from n = 6 new participants scanned for this study). As a result, there were three versions of the component localizer, differing in the minor details of stimulus presentation and scanning parameters (see **Supplemental Table 1**), but which otherwise were very similar. All three versions contained the same set of 30 2-second natural sounds. In all versions, stimuli were presented during scanning in a “mini-block design,” in which each 2-second sound was repeated multiple times in a row. For version 1, stimuli were repeated 5 times in a row, and 3 times for versions 2 and 3. During scanning, stimuli were presented over MR-compatible earphones (Sensimetrics S14) at 75 dB SPL. Each 2-second stimulus was presented in silence, with a single fMRI volume collected after each presentation (i.e., “sparse scanning”; Hall et al., 1999). A short ∼200 ms buffer period was included between the stimuli and the volume acquisition, leading to a TR of approximately 3.4 seconds and a TA of approximately 1 second (exact scanning parameters for each version of the localizer can be found in **Supplemental Table 1**). To encourage participants to pay attention to the sounds, either the second or third repetition in each “mini-block” was 8dB quieter (presented at 67 dB SPL), and participants were instructed to press a button when they heard this quieter sound. Data pre-processing, denoising, and initial GLM analyses were the same as in the main experiment (see “*fMRI data acquisition and analysis*” below).

Despite the fact that participants were run on different versions of the component localizer experiment, we found that the components from Norman-Haignere et al. (2015) explain similar amounts of variance for each group of participants (n = 2 from Norman-Haignere et al. 2015, median R^2^ = 0.78, SD = 0.04; n = 7 from Boebinger et al. 2021, median R^2^ = 0.80, SD = 0.04; n = 6 new participants, median R^2^ = 0.83, SD = 0.04; small effect of group, ɛ^2^ = 0.03).

#### Component weight estimation

To estimate each participant’s component weights (𝑊), we multiplied each participant’s data matrix of voxel responses to the 30 natural sound stimuli (𝐷_)*_; sound x voxel) by the pseudoinverse of the known component response matrix for those same 30 sounds (𝑅_)*_; sound x component):

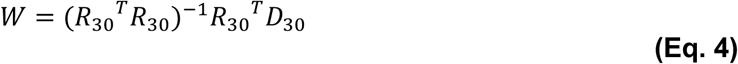

Multiplying by the pseudoinverse is equivalent to performing ordinary least-squares regression. For visualization purposes, component weights were averaged across participants in standardized anatomical coordinates (Freesurfer’s FsAverage template), and plotted with a color scale that spans the central 95% of the weight distribution for each component.

#### Component response estimation

The goal of the current study was to then use participants’ inferred component weights to estimate the component responses to each of the stimulus conditions in the main experiment. To do this, we multiplied participants’ data matrix of voxel responses to the experimental stimuli (𝐷_+,-_) by the pseudoinverse of the estimated component weights:

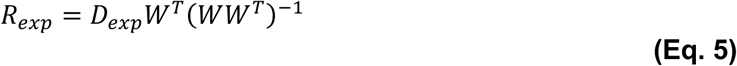

Unless otherwise noted in the text, we included all voxels in a large anatomical mask region encompassing bilateral superior temporal and posterior parietal cortex (shown as colored regions in the figures). For analyses comparing different cortical regions (e.g., right vs. left hemisphere, anterior vs. posterior auditory cortex), we estimated the component response for each cortical region separately, using the voxel responses and estimated component weights from the corresponding subset of voxels.

Because the component responses are estimated as a weighted combination of responses across voxels – with each voxel contributing in proportion to its component weight – the spatial specificity of an estimated component response is determined by the localization of its weight map. For example, since the music component’s weights are concentrated in anterior and posterior auditory cortex, its estimated response is driven primarily by voxels in those regions rather than by voxels distributed uniformly across the anatomical mask. Multiplying by the pseudoinverse of the weight matrix rather than the weights directly corrects for correlations between overlapping components, but does not change this basic interpretation. When we repeat our analyses separately in anterior and posterior temporal cortex, we are therefore effectively isolating the component response within the local voxel populations that contribute most strongly to it.

### Main experiment

#### Stimuli

The stimuli for the main experiment consisted of 10-second clips of intact and scrambled versions of synthetic music, as well as several non-music control conditions. Specifically, we created versions of each music clip in which we manipulated (1) the note pitches, (2) the note onset times, and (3) both the note pitches and onset times. So that we could examine the effect of manipulating temporal structure in the absence of pitch structure, we also included a set of synthetic drum stimuli and scrambled versions of each of those stimuli in which we manipulated the note onset times.

To construct these stimuli, we first selected a set of 20 music clips from the stimulus set used in a previous study by our group (Fedorenko et al., 2012). These clips originated from polyphonic musical pieces from a variety of genres common in modern Western musical traditions, each with complex tonal and rhythmic content, and were rendered in MIDI (musical instrument digital interface) format. Each piece contained multiple instrumental tracks, which were all converted to a piano timbre for use in the experiment. The keys of the clips were assigned by drawing (without replacement) from a pool of all 12 major and 12 minor keys (preserving the mode, i.e. “major” vs. “minor”), and the key of each clip was then changed by adding/subtracting a constant (corresponding to the difference in semitones between the original and desired keys) to the note pitches. As a result, the 14 major clips included in the stimulus set spanned all 12 major keys (with C# and G# major occurring twice), and the 6 minor clips were all in different keys.

We then created three additional versions of each music clip (20 unique clips x [1 intact + 3 scrambled versions] = a total of 80 MIDI stimuli), in which the musical structure was manipulated by scrambling either the pitches and/or timing of the notes (**Figure 1**). MIDI represents music as a matrix of notes, with different columns specifying the onset, duration, pitch, and instrumentation of the notes, which makes it possible to manipulate these features independently and to scramble across-note structure without altering the acoustic properties of individual notes. In all cases, within-chord structure was preserved by performing the scrambling operation on clusters of notes with a similar onset time (defined for our purposes as within 0.1 beats of each other), rather than on individual notes (**Figure 1A** & **B**). The tempi of the MIDI stimuli ranged from 72 to 220 bpm, so a note cluster threshold of 0.1 beats corresponded to note onsets that occur within a time interval that varied between 27 and 83ms, depending on the tempo. This is a more conservative scrambling procedure than was used in our previous study using related methods (Fedorenko et al., 2012). For the pitch-scrambled music clips, each note cluster was shifted either up or down by 0-2 semitones (sampled from a discrete, uniform distribution), which disrupted both melodic and tonal structure by abolishing the stereotyped profile of pitches within a key and thus the notion of a “tonal center” (**Figure 1C** & **D**). For time-scrambled music clips, the inter-onset intervals between note clusters were jittered by adding an amount drawn from a uniform distribution spanning ±0.25 beats (with the range of tempi of our stimuli, 0.25 beats corresponded to between 68-208ms), but with the constraint that no inter-onset interval could be shorter than 0.1 beats. To ensure that the total duration of the time-scrambled clip was the same as the original version, we divided the vector of jittered inter-onset intervals by its mean and then multiplied it by the mean of the original vector of inter-onset intervals. Next, the jittered inter-onset intervals were randomly reordered, and the note onset times of the scrambled piece were derived from these reordered inter-onset intervals (via cumulative summation). This process effectively eliminated any isochronous beat or metrical hierarchy (**Figure 1E**). For the music clips that were both pitch- and time-scrambled, they first went through the time-scrambling procedure and then the pitch-scrambling procedure. This set of 80 MIDI stimuli was supplemented by a set of 20 MIDI drum stimuli, chosen to represent a variety of genres, with all percussion events assigned to the standard General MIDI drum kit patch. We also created versions of these drum stimuli that were time-scrambled following the procedure described above (**Figure 1F**), for a total of 40 MIDI drum stimuli. Each MIDI file was converted into audio using AppleScript and QuickTime Player 7 (Ellis, 2014).

**Figure 1.**
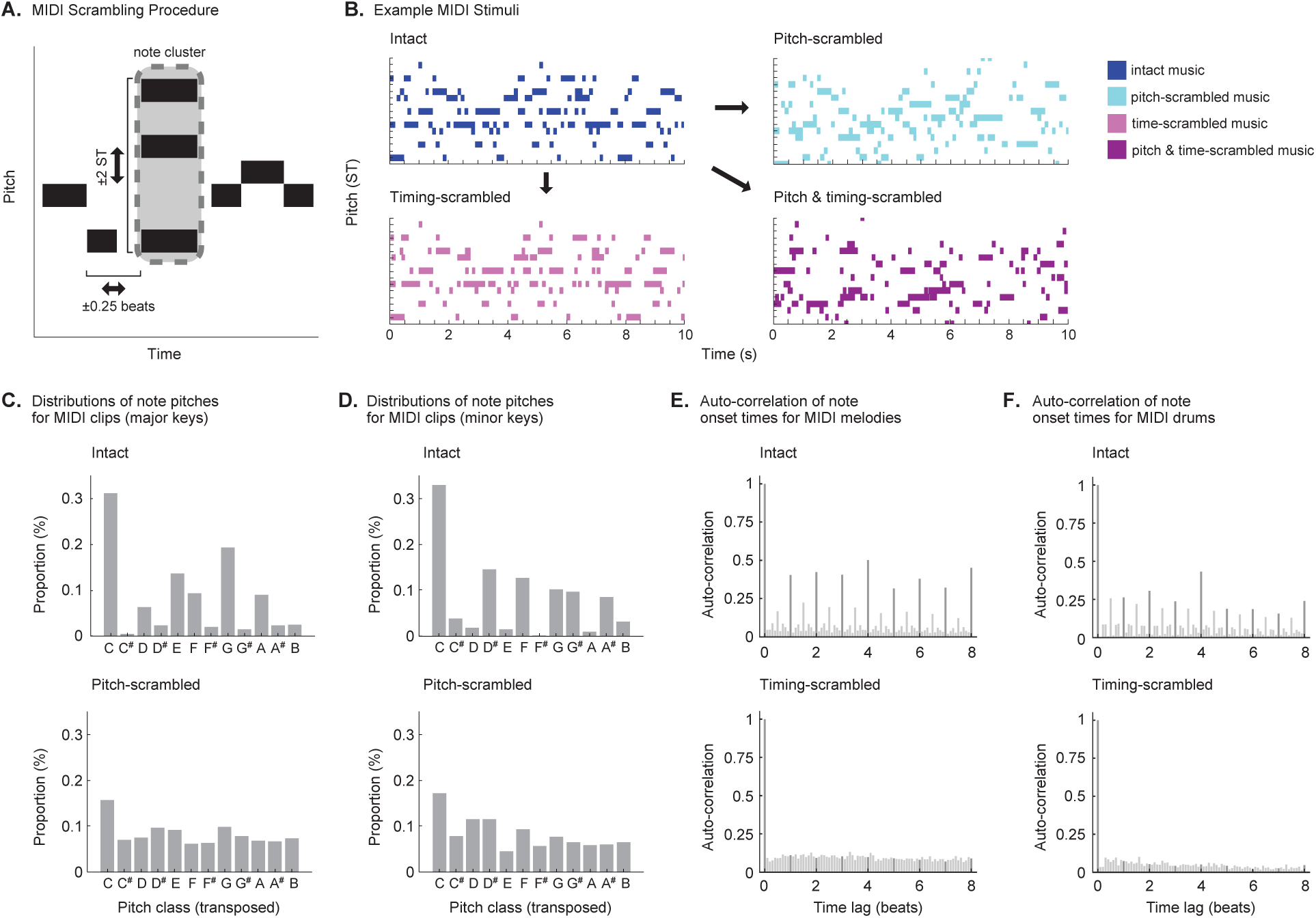
Scrambling procedure for MIDI music stimuli. **A.** Schematic of scrambling procedure. First, notes with the same onset (within 0.1 beats) were grouped together and scrambled as a unit. For pitch-scrambling, each note cluster was shifted by up to ±2 semitones. For timing-scrambling, the inter-onset intervals (IOIs) between note clusters were jittered by adding an amount drawn from a uniform distribution spanning ±0.25 beats (with the constraint that no IOI could be shorter than 0.1 beats). The jittered IOIs were normalized to ensure the total duration of the timing-scrambled music clip was the same as the original intact version and were randomly reordered. For the music clips that were both pitch- and timing-scrambled, they first went through the timing-scrambling procedure and then the pitch-scrambling procedure. **B.** Intact and scrambled versions of an example MIDI stimulus. **C.** Histograms showing the distribution of note pitch classes averaged across the 14 MIDI clips in major keys, in both their intact (top) and pitch-scrambled (bottom) versions. For the purpose of this analysis, all clips were transposed to C major. Pitch class distributions were weighted by note durations, as computed using the *“pcdist1”* function in the MATLAB MIDI Toolbox (Eerola & Toiviainen, 2004). For the intact music clips (top), certain pitch classes are more common than others, reflecting the tonal hierarchy typical of major keys in Western music. The pitch-scrambling procedure (bottom) disrupts both melodic and tonal structure, leveling out the distribution of pitch classes to a certain extent. **D.** Same as **C**, but for the 6 MIDI clips in minor keys. **E.** Histograms showing the autocorrelation of note onset times in units of beats averaged across all 20 intact (top) MIDI stimuli and their timing-scrambled versions (bottom). Onset times were weighted by note durations, as computed using the *“onsetacorr”* function in the MATLAB MIDI Toolbox (Eerola & Toiviainen, 2004). For the intact music clips (top), peaks can be seen on every beat (darker gray bars) that reflect the isochronous beat and metrical structure typical of Western music. The timing-scrambling procedure (bottom) disrupts this metrical structure, resulting in no clear peaks in the autocorrelation function at multiples of the beat. **F.** Same as **E**, but for the 20 MIDI drum stimuli.

To complement these 120 MIDI music stimuli, we selected 16 additional non-music stimuli to serve as controls for assessing selectivity. Note that we use “selectivity” not to imply a binary distinction between responses to “music” and “non-music” stimuli, but instead to allow for a graded pattern of responses in which the response magnitude to the preferred stimulus category is greater than to all other tested categories (Downing et al., 2007; Fedorenko et al., 2010; Leaver & Rauschecker, 2010; Op De Beeck et al., 2008). Specifically, we chose 4 English speech excerpts (taken from audiobook recordings and radio broadcasts), 4 foreign speech excerpts (German, Hindi, Italian, Russian), 4 animal vocalizations (chimpanzees, cow, sheep, wolf), and 4 environmental sounds (footsteps, ping-pong ball, power saw, copy machine). All stimuli were resampled to 20 kHz, trimmed to be 10 seconds long, windowed with a 25 ms half-Hanning-window ramp, and normalized to have the same RMS.

Because the MIDI scrambling manipulations were intended to preserve some surface aspects of musical structure, an online experiment (via Amazon’s Mechanical Turk) was used to ensure that the manipulation produced detectable changes in musical structure. In this experiment, participants listened to the same 120 MIDI stimuli used in the full experiment (n = 46; 16 self-identified female, 27 self-identified male, 0 self-identified non-binary, and 3 did not answer; mean age = 42.9 years, SD = 11.9 years; an additional 7 participants completed the experiment but were excluded due to performance of <60% in every condition). Stimuli were presented in pairs, with each trial containing an intact and scrambled version of the same MIDI stimulus. Participants were instructed that one of the stimuli was “corrupted by having the melody and/or the note timing disrupted,” and were told to choose which of the two music clips had been “corrupted.” Stimuli were blocked by condition, so that each participant completed four blocks of 20 trials each (i.e., intact vs. pitch-scrambled, intact vs. time-scrambled, intact vs. pitch-and-time-scrambled, and intact vs. time-scrambled drums); participants were told which type of “corruption” to listen for in each block. Prior to completing the experiment participants completed a headphone check to help ensure they were wearing headphones or earphones (Woods et al., 2017); we have found that this headphone check is typically sufficient to obtain performance on par with that of experiments conducted in the lab under more controlled listening conditions (McPherson et al., 2020; Traer et al., 2021; Woods & McDermott, 2018).

Performance was well above chance in all four blocks, indicating that the scrambling manipulation was clearly detectable in all conditions (pitch-scrambling: median = 87.5%, interquartile range or IQR = 20%; time-scrambling: median = 100%, IQR = 5%; pitch-and-time-scrambling: median = 100%, IQR = 5%; time-scrambling drums: median = 100%, IQR = 5%; one-sample Wilcoxon signed rank tests comparing percent correct to chance performance of 50%; all p’s < 1e-08, two-tailed).

#### Stimulus presentation and scanning procedure

During the scanning session for the main experiment, auditory stimuli were presented over MR-compatible earphones (Sensimetrics S14) at 80 dB SPL. The 10-second stimuli were presented back-to-back with a 500 ms inter-stimulus interval. To encourage participants to pay attention to the stimuli, each stimulus either ramped up (by 6 dB) or down (by 9 dB) in level over 1 second starting at the 5s point, and the participant indicated whether each stimulus got “louder” or “quieter” via button press. The increment and decrement values were chosen to approximately equate the subjective salience of the level change. Participants’ average performance on this task was 97.1% correct (SD = 2%), and performance never fell below 85% correct for any participant on any run.

The scanning session for the main experiment consisted of twelve 7.3-minute runs (all participants completed 12 runs, except for one participant who only completed 8 runs), with each run consisting of thirty-four 10-second stimuli and six 10-second silent periods during which no sound was presented. These silent blocks were the same duration as the stimuli and were distributed randomly throughout each run, providing a baseline. The thirty-four stimuli presented in each run consisted of five stimuli from of each of the MIDI music conditions (“intact,” “pitch-scrambled,” “time-scrambled,” “pitch-and-time-scrambled,” “intact drums,” and “time-scrambled drums”), and one stimulus each from the four non-music control conditions (“English speech,” “foreign speech,” “animal sounds,” “environmental sounds”). To maximize the temporal interval between a given MIDI music stimulus and its scrambled versions, only one version of a given MIDI stimulus (i.e. intact, pitch-scrambled, time-scrambled, pitch-and-time-scrambled) was presented per run for all but three participants (the remaining three participants were run on an earlier version of the experiment that was identical except without the constraint maximizing the temporal interval between intact and scrambled versions of a given MIDI stimulus, and with an extra null block in each run). The full set of 136 stimuli was repeated every four runs, meaning that each individual stimulus was presented a total of three times during the course of the 12 experimental runs (twice for the participant who only completed 8 runs). The entire scanning session lasted approximately 2 hours.

#### fMRI data acquisition and analysis

MRI data were collected at the Athinoula A. Martinos Imaging Center of the McGovern Institute for Brain Research at MIT, on a 3T Siemens Trio with a 32-channel head coil. Because the sound stimuli were long (10 seconds), we used continuous instead of sparse scanning (TR = 2.1 sec, TE = 30ms, 90-degree flip angle, 3 discarded initial acquisitions). Each functional acquisition consisted of 46 roughly axial slices (oriented parallel to the anterior-posterior commissure line) covering the whole brain, with voxel size 2 x 2 x 2.8mm (96 x 96 matrix, 10% slice gap). A simultaneous multi-slice (SMS) acceleration factor of 2 was used in order to minimize acquisition time. To localize functional activity, a high-resolution anatomical T1-weighted MPRAGE image was obtained for each participant (1 mm isotropic voxels).

Preprocessing and data analysis were performed using FSL software and custom Matlab scripts. Functional volumes were motion-corrected, slice-time-corrected, skull-stripped, linearly detrended, and aligned to each participant’s anatomical image (using FLIRT and BBRegister; (Greve & Fischl, 2009; Jenkinson & Smith, 2001). Motion correction and functional-to-anatomical registration was done separately for each run. Each participant’s preprocessed data were then resampled to the cortical surface reconstruction computed by FreeSurfer (Dale et al., 1999), registered to the FsAverage template brain, and then smoothed on the surface using a 3mm FWHM kernel to improve the signal-to-noise ratio (SNR) of the data by removing local, uncorrelated noise.

GLM-denoise (Kay et al., 2013) was used to further improve SNR and then estimate the response to the stimuli using a general linear model (GLM). A separate boxcar regressor was used for each of the 10 experimental conditions, which were convolved with a hemodynamic response function (HRF) that specifies the shape of the BOLD response. GLM-denoise assumes a common HRF that is shared across all conditions and voxels, the shape of which is estimated from the data using an iterative linear fitting procedure (Kay et al., 2013).

To remove large-scale correlated noise across voxels, GLM-denoise also includes noise regressors based on principal components analysis (PCA). These PCA-based regressors were derived from the time series of a “noise pool” consisting of voxels with <0% of variance explained by the task regressors (R^2^ was measured using leave-one-run-out cross-validation, which is negative when the null hypothesis that all beta weights equal 0 outperforms the model on left-out data). The optimal number of noise regressors was determined by systematically varying the number of principal components included in the model, and estimating the R^2^ of each model using leave-one-run-out cross-validation.

GLM-denoise then performs a final fit of the model, and estimates of beta weight variability are obtained by bootstrapping across runs 100 times. The final model estimates are taken as the median across bootstrap samples, and these beta weights were converted to units of percent BOLD signal change by dividing by the mean signal intensity in each voxel. GLM-denoise was applied to the data from all runs for each participant. The resulting beta weights were downsampled to a 2mm isotropic grid on the Freesurfer-flattened cortical surface. To accommodate the partial-brain coverage for two of the participants (see “version 3” in **Supplemental Table 1**), we limited all analyses to voxels within a large anatomical mask region encompassing bilateral superior temporal and posterior parietal cortex (shown as colored regions in the figures). All subsequent analyses were conducted in this downsampled 2D surface space, and for ease of description, we refer to the elements as “voxels” throughout the paper.

### Anterior and posterior auditory cortex anatomical regions of interest (ROIs)

Our previous work has found that the music-selective component weights are clustered into “hotspots,” one in anterior STG and the other in posterior STG (**Figure 2A**). To test whether these two clusters might represent different aspects of musical structure, we carefully selected a dividing point that best subdivided the anterior and posterior clusters in individual participants. We did this by drawing a straight line on the downsampled 2D cortical surface, approximately halfway along Heschl’s gyrus (HG) and roughly perpendicular to STG (see white dashed line in **Figure 2B**). Importantly, we do not simply average the response of all the voxels in these ROIs. Instead, our component analysis finds a weighted combination of the voxels within each ROI that isolates music-selective populations. Thus, what is critical is that the dividing line optimally separates the anterior and posterior music-selective neural populations.

**Figure 2.**
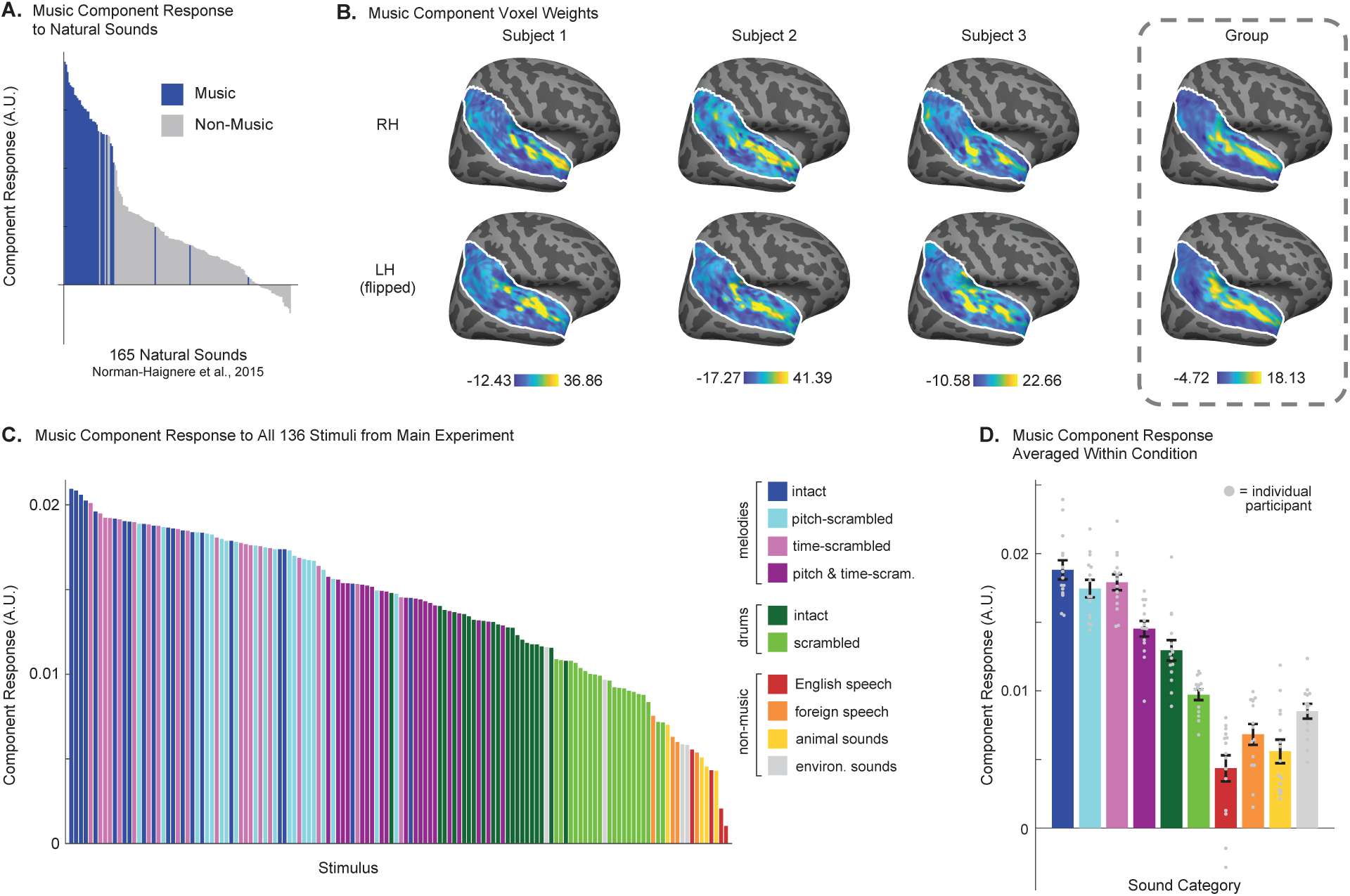
Music component response to MIDI music and non-music control stimuli. **A.** Music component response profile across all 165 natural sounds from Norman-Haignere et al. (2015) study. Sounds categorized as both “instrumental music” and “vocal music” (as determined by raters on Amazon Mechanical Turk) are shaded blue to highlight the degree of music selectivity. **B.** Spatial distribution of music component voxel weights for participants in the current study. Shown for three example participants transformed to standardized anatomical coordinates (FreeSurfer’s FsAverage template), and averaged across participants (n = 15). Color scale spans the central 95% of the weight distribution. **C.** Music component response to each of the 136 individual stimuli in the main experiment, averaged across participants. **D.** Music component response to the stimulus conditions in the main experiment, averaged across participants. Each gray dot corresponds to one participant’s component response to a given condition. Scatter plots are within-subject, such that differences between participants are removed by subtracting each participant’s mean response across conditions and then adding the global mean across participants. Error bars indicate ±1 within-subject SEM.

### Statistics

To compare responses across two conditions, such as intact vs. scrambled drums, two-sided Wilcoxon signed-rank tests were used and effect sizes were calculated as:

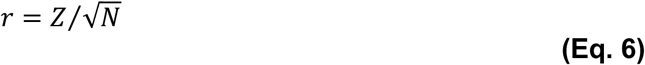

where Z is the z-score output of the Wilcoxon signed rank test, and N is the number of pairs (i.e., the number of participants). We note that several Wilcoxon signed-rank tests yielded identical Z statistics, p-values, and effect sizes because the same number of paired observations was used in each contrast and all paired differences were in the same direction with no ties; under these conditions, rank-based non-parametric tests reach their maximal statistic and produce identical results regardless of difference magnitude.

We used repeated-measures ANOVAs for comparisons across more than two conditions or factors. In a 2 x 2 within-subjects design with complete data, this approach is equivalent to fitting a linear mixed-effects (LME) model with random intercepts for participants.

In all cases, we used the Shapiro-Wilk test to verify that the standardized residuals for every combination of factors were normally distributed, and the results of this test were non-significant (p > 0.05) unless otherwise noted in the text. When the assumption of normality was not met, the significance of the F-statistic was evaluated using approximate permutation tests. To do this, we randomized the assignment of the data points for each participant across the conditions being tested 10,000 times, recalculated the F-statistic for each permuted sample, and then compared the observed F-statistic to this null distribution to determine significance. Specifically, permutations were restricted such that only the structure relevant to the effect under test was disrupted, while the structure of all other factors was preserved. For main effects, the condition labels for the factor of interest were shuffled across its two levels within each participant, independently of all other factors. For interaction effects, condition labels were swapped between levels of one factor within each level of the other factor(s), preserving the marginal distributions of all main effects while disrupting only the contingency between factors that gives rise to the interaction. The ‡ symbol in Tables 1 and 2 identifies the tests reported in those tables for which the permutation approach was used. We also explicitly note in the text where permutation tests were used for ANOVAs that do not appear in the tables. Note that there was no need to check for violations of sphericity, because sphericity necessarily holds for repeated-measures factors with only two levels. For both parametric and non-parametric ANOVAs, effect sizes were quantified using partial eta-squared (η_p_^2^). ANOVAs were conducted across participants (n = 15).

**Table 1.** ANOVA results for all auditory cortical response components. ‡ = Shapiro-Wilk test indicated that the standardized residuals for one or more combinations of factors were non-normal (p < 0.05), so the significance of the F-statistic was evaluated using a non-parametric permutation test randomizing the assignment of the data points across the relevant conditions 10,000 times (see “*Statistics*” section of Methods). * = Significant at p < 0.05, ** = significant at p < 0.01, *** = significant at p < 0.001.

|  |  | Low freq. | High freq. | Temp. mod. | Pitch | Speech | Music |
| --- | --- | --- | --- | --- | --- | --- | --- |
| Pitch-scrambling main effect | F-value | 1.93 | 12.88** | 19.75*** | 0.85 | 45.53*** | 39.50*** |
|  | p-value | 0.19 | 0.003 | 2.0e-04‡ | 0.37 | 9.36e-06 | 2.01e-05 |
| | $\eta_p^2$ | 0.12 | 0.5 | 0.59 | 0.06 | 0.76 | 0.74 |
| Timing-scrambling main effect | F-value | 0.66 | 53.96*** | 20.42** | 1.67 | 0.18 | 15.20** |
|  | p-value | 0.43 | 3.60e-06 | 0.006‡ | 0.22 | 0.68 | 0.002 |
| | $\eta_p^2$ | 0.05 | 0.79 | 0.42 | 0.11 | 0.01 | 0.52 |
| Pitch x timing interaction | F-value | 0.69 | 6.85 | 2.06 | 4.88 | 32.11*** | 26.91*** |
|  | p-value | 0.42 | 0.02 | 0.18‡ | 0.05 | 5.80e-05 | 1.38e-04 |
| | $\eta_p^2$ | 0.05 | 0.133 | 0.13 | 0.26 | 0.7 | 0.66 |

**Table 2.**
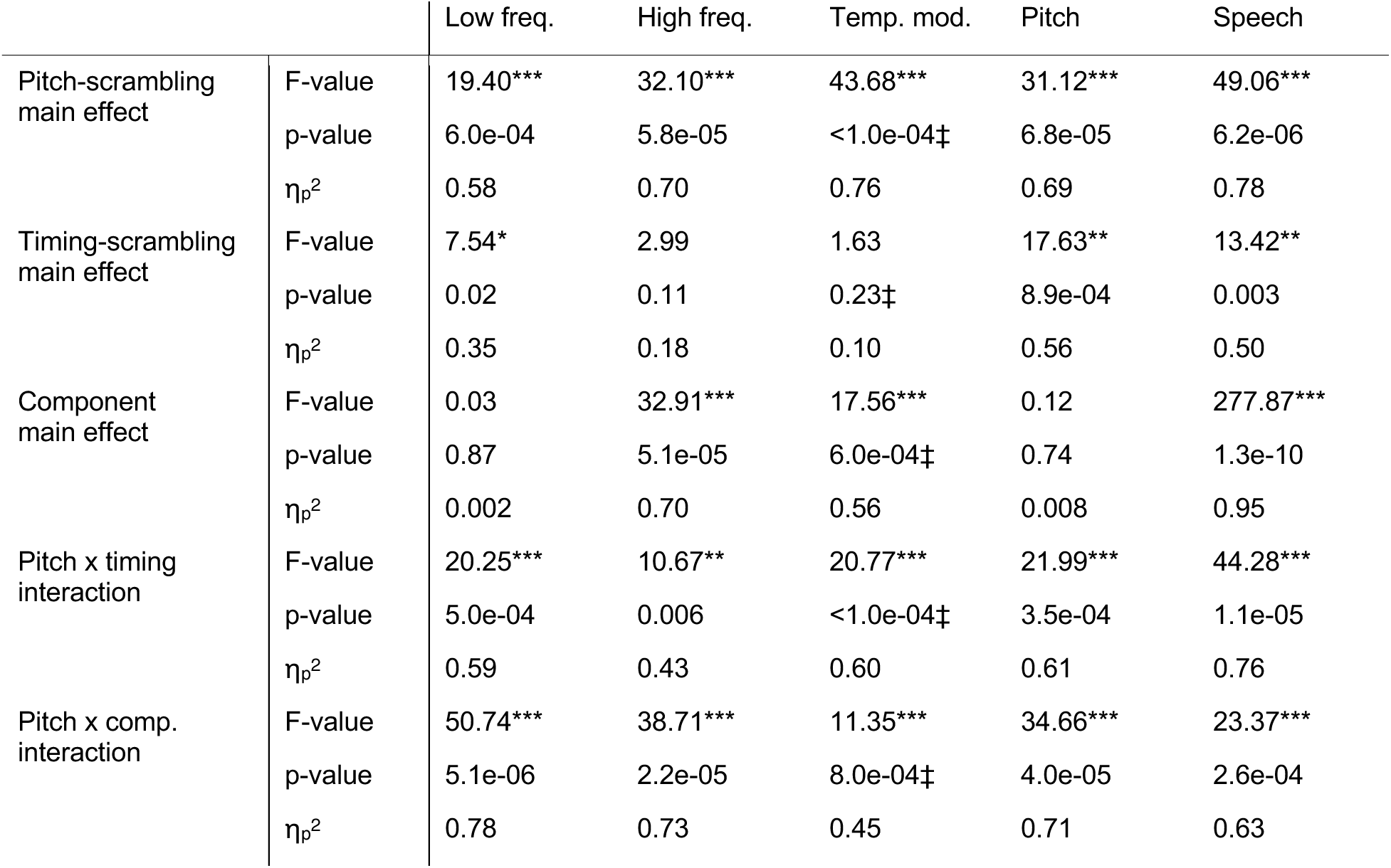
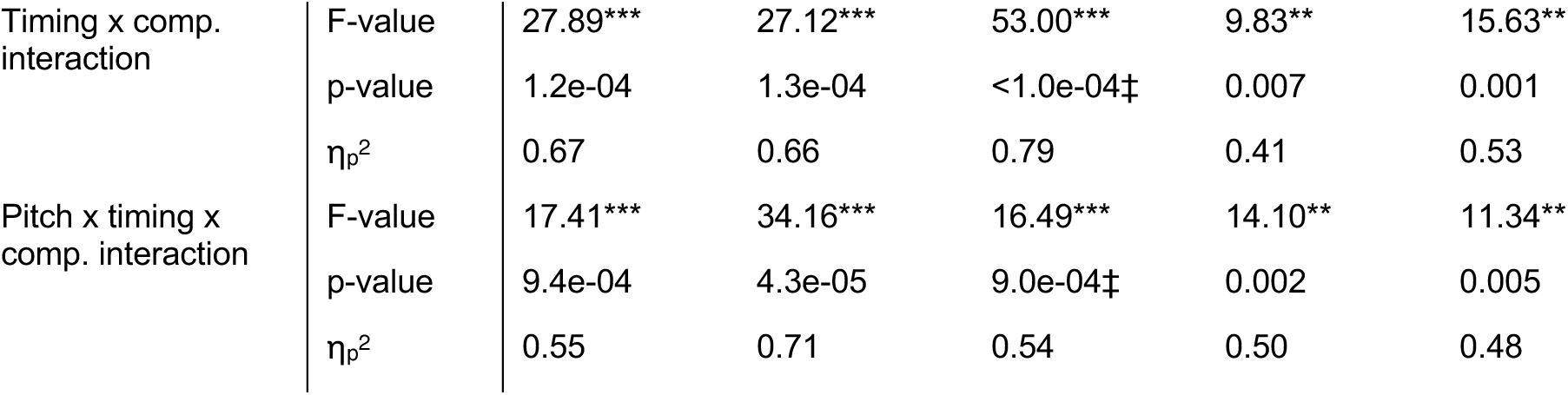
Results for 3-way repeated-measures ANOVAs comparing the music component to each of the other auditory cortical response components. ‡ = Shapiro-Wilk test indicated that the standardized residuals for one or more combinations of factors were non-normal (p < 0.05), so the significance of the F-statistic was evaluated using a non-parametric permutation test randomizing the assignment of the data points across the relevant conditions 10,000 times (see “*Statistics*” section of Methods). * = Significant at p < 0.05, ** = significant at p < 0.01, *** = significant at p < 0.001.

For all figures, we used “within-subject” scatter plots and error bars (Loftus & Masson, 1994), which remove between-subject variance by subtracting each participant’s mean response across conditions. The global mean across participants is then added to this de-meaned data before plotting and computing the standard error (SEM).

## RESULTS

### Music-selective neural populations are sensitive to the patterning of musical notes

We sought to test whether music-selective neural populations in human non-primary auditory cortex are driven by the patterning of notes in pitch or time. We therefore measured responses to MIDI music and drum stimuli that had been scrambled in pitch or time from 15 participants. Because music selectivity is weak when measured using standard voxel-wise fMRI analyses, we utilized techniques from our prior work (Norman-Haignere et al., 2015) to isolate music-selective responses from other, potentially spatially overlapping brain responses (**Figure 2A**). We used our component localizer data to estimate a set of voxel weights (**Figure 2B**) for each participant that best approximated these previously inferred component response profiles. The weights for the music component showed at least two distinct clusters in anterior and posterior non-primary auditory cortex, replicating prior studies. These clusters correspond to the voxels that contribute most strongly to the estimated music-component responses. We then used these weights to infer the component responses to the stimuli in the main experiment (**Figure 2C & D**) (by multiplying the pseudoinverse of the weights with the participant’s voxel responses from the main experiment). Because voxels with larger weights contribute more strongly to the estimated component response, the spatial clustering of weights means that these responses are driven primarily by activity in the anterior and posterior clusters rather than uniformly across temporal cortex. As expected, the component replicated its selectivity for music, responding most strongly to intact music, and minimally to the non-music control stimuli (**Figure 2C & D**). Further, the response of the music component to drum stimuli was significantly higher than the non-music sounds (Wilcoxon signed-rank test, *Z* = 3.35, p = 8.05e-04, r = 0.87) but significantly lower (*Z* = 3.35, p = 8.05e-04, r = 0.87) than the pitched music stimuli, replicating previous findings (Boebinger et al., 2021).

Our first question was the extent to which the music-selective component’s response is driven by selectivity for note-pattern structure (i.e., the organization of notes into coherent pitch or temporal sequences) vs. note-level structure (i.e., the acoustic properties of individual notes). These two accounts make different predictions about the pattern of responses across intact music, fully scrambled music, and non-music stimuli. Selectivity for note-pattern structure predicts a stronger response to intact compared with fully scrambled music, since scrambling disrupts these note patterns. Selectivity for the acoustic properties of individual notes, by comparison, predicts that even fully scrambled music should evoke reliably stronger responses than all other non-music stimuli, since scrambled music preserves note-level acoustic structure that is absent in sounds from other categories.

We therefore evaluated this hypothesis using pairwise statistical tests between the responses to intact music, pitch-and-time-scrambled music, and non-music control stimuli. Strikingly, we found that for all 20 of the MIDI clips tested, the response to the intact version was stronger than the response to the corresponding fully scrambled version (**Figure 2C**), resulting in an effect that was highly significant (*Z* = 3.41, p = 6.55e-04, r = 0.88). On average, the magnitude of the response to pitch and time-scrambled stimuli was approximately 23% lower than that for fully intact music, suggesting that note-pattern structure makes a non-trivial contribution to this component’s selectivity for music.

At the same time, the response to fully scrambled music stimuli was still substantially higher than that for non-music stimuli, with the response to every single pitch-and-time-scrambled stimulus being higher than the response to every other non-music stimulus tested. On average, the response to non-music stimuli was only 56% as strong as the response to pitch-and-time-scrambled music, and this highly significant difference (*Z* = 3.35, p = 8.05e-04, r = 0.87) is inconsistent with a purely sequence-level account and instead indicates sensitivity to note-level structure even when higher-order note patterns are disrupted. Consistent with this interpretation, the 23% drop in the response of the music-selective component for pitch-and-time-scrambled was less than that observed in prior studies for manipulations that disrupt note structure (e.g., with quilting or modulation-matched sounds) but preserve other low-level acoustic properties (Norman-Haignere et al., 2015; Norman-Haignere & McDermott, 2018).

#### Music-selective neural populations are sensitive to note patterns in both pitch and time

Does sensitivity to the patterning of notes reflect sensitivity to pitch patterns, time patterns, or both? It is impossible to perfectly isolate pitch from temporal structure in complex polyphonic music. For that reason, our drum stimuli provide the clearest test of temporal structure, since they lack definite pitch – they may sound “high” or “low,” but do not have an identifiable pitch. Even though responses to drum stimuli were lower overall than to pitched music, we observed a robust scrambling effect (*Z* = 3.41, p = 6.55e-04, r = 0.88), with a stronger response to intact vs. scrambled versions for all 20 of the drum stimuli tested. On the other hand, our most direct manipulation of pitch structure is the difference between time-scrambled and pitch-and-time scrambled conditions, both of which lack coherent rhythmic structure. This difference is highly significant (Z = 3.41, p = 6.55e-04, r = 0.88) with a stronger response to each of the 20 time-scrambled stimuli than to their pitch-and-time-scrambled counterparts derived from the same original melody. Together, these results suggest that music-selective neural populations are sensitive to both pitch and temporal patterns in music.

Because our scrambling manipulation was designed so that each music clip appeared in all four conditions (intact, pitch-scrambled, time-scrambled, and pitch-and-time-scrambled), we used a repeated-measures ANOVA to examine the effect of disrupting pitch and temporal structure. We observe clear main effects of both pitch and time scrambling (pitch: F(1, 14) = 39.50, p = 2.01e-05, η_p_^2^ = 0.74; time: F(1, 14) = 15.20, p = 0.002, η_p_^2^ = 0.52), as well as a significant interaction between pitch and temporal structure. The interaction was driven by the fact that scrambling in one domain (pitch or time) was larger if the other domain was scrambled as well (F(1, 14) = 26.91, p < 1.38e-04, η_p_^2^ = 0.66). In fact, the responses to the pitch-scrambled and time-scrambled conditions are only somewhat weaker than the response to the intact condition, with the pairwise comparison between intact and time-scrambled conditions not reaching statistical significance (Z = 1.48, p = 0.14, r = 0.38; intact vs. pitch-scrambled, Z = 2.78, p = 0.005, r = 0.72). This pattern suggests that the presence of either pitch or temporal structure substantially preserves the music component’s response, consistent with partial note-pattern structure being sufficient to drive a strong response. This interpretation is grounded primarily in the significant interaction, which indicates that the effect of scrambling in one domain is considerably reduced when the other domain remains intact, and is further supported by our drum scrambling results, where the presence of coherent temporal structure produces a stronger response even when the stimuli lack stable note-level pitch.

### Sensitivity to temporal and pitch patterns is widely distributed across anterior and posterior music-selective neural populations in both the left and right hemispheres

Music-selective neural populations are concentrated in at least two widely spaced clusters in anterior and posterior non-primary auditory cortex. Given this finding, and prior studies suggesting differences in spectrotemporal modulation tuning in anterior vs. posterior auditory cortex (Hamilton et al., 2018; Hullett et al., 2016b; Norman-Haignere et al., 2015; Santoro et al., 2014b; Schönwiesner & Zatorre, 2009), we next investigated whether there was a clear difference in sensitivity to musical structure between anterior and posterior regions. We divided auditory cortex into anterior and posterior regions of interest (ROIs) in a way that best separated the anterior and posterior clusters in individual participants, approximately halfway along Heschl’s gyrus (HG) and roughly perpendicular to STG (see white dashed line in **Figure 3A**), and used our component analysis to find a weighted combination of the voxels within each ROI that isolated music-selective neural populations. We also sought to test whether there might be differences in sensitivity for pitch pattern and temporal pattern structure between the two hemispheres, motivated by prior hypotheses about hemispheric organization (Albouy et al., 2020; Zatorre & Belin, 2001).

**Figure 3.**
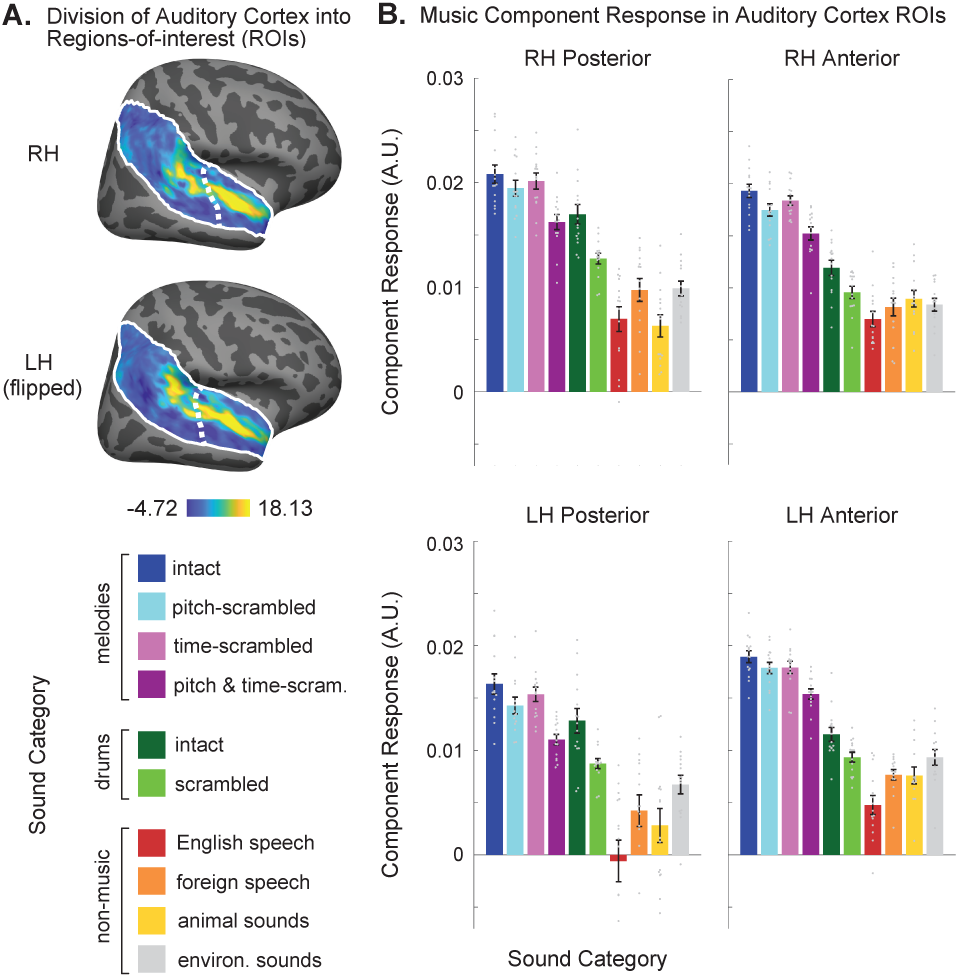
**A.** Spatial distribution of music component voxel weights, averaged across participants in standardized anatomical coordinates (FreeSurfer’s FsAverage template). Color scale spans the central 95% of the weight distribution. The white dashed line shows the dividing line between anterior and posterior ROIs. This line was defined by drawing a straight line on the downsampled 2D cortical surface to maximize the separation of individual participants’ clusters of music selectivity. **B.** Music component response to the stimulus conditions in the main experiment, plotted separately for the anterior and posterior ROIs in the left and right hemispheres. Each gray dot corresponds to one participant’s component response to a given condition. Scatter plots are within-subject, such that differences between participants are removed by subtracting each participant’s mean response across conditions and then adding the global mean across participants. Error bars indicate ±1 within-subject SEM.

To address this question, we repeated our entire analysis using component weights and voxel responses from each combination of hemisphere and ROI separately (i.e., right hemisphere anterior, right hemisphere posterior, left hemisphere anterior, and left hemisphere; **Figure 3A**). Overall, we found that the response pattern across conditions was strikingly similar for all four subdivisions (**Figure 3B**). A 4-way repeated-measures ANOVA incorporating both hemisphere and ROI as factors showed significant main effects of pitch and temporal scrambling and their interaction (pitch-scrambling: F(1,14) = 40.55, p = 1.75e-05, η_p_^2^ = 0.74; time-scrambling: F(1,14) = 16.63, p = 0.001, η_p_^2^ = 0.54; interaction: F(1,14) = 26.54, p =1.47e-04). Importantly, we did not observe any significant interactions between hemisphere and either type of scrambling manipulation (pitch-scrambling or time-scrambling) or between ROI and either type of scrambling manipulation (all p’s > 0.07). We did observe a significant main effect of hemisphere (F(1,14) = 18.21, p = 7.81e-04, η_p_^2^ = 0.57) and a significant ROI x hemisphere interaction (F(1,14) = 18.47, p = 7.37e-04, η_p_^2^ = 0.57), but these are driven by differences in overall response magnitudes across all conditions (as can be seen in **Figure 3B**), which is not particularly meaningful because response magnitude is influenced by a myriad of factors, including non-neural factors (e.g., vascularization).

To more directly test the sensitivity to temporal scrambling across auditory cortex, we ran a second repeated-measures ANOVA on the response to the intact and scrambled drum conditions. We found a significant effect of temporal scrambling (F(1,14) = 45.87, p < 0.001, η_p_^2^ = 0.77; due to non-normality the significance of the F-statistics involving drum conditions were all determined using non-parametric permutation tests, see “Statistics” section of Methods for more details), with both anterior and posterior ROIs showing significant effects of drum scrambling (anterior ROI: Z = 3.35, p = 08.1e-04, r = 0.87; posterior ROI: Z = 3.41, p = 6.5e-04, r = 0.88). The scrambling effect was larger in the posterior ROI (scrambling x ROI interaction: F(1,14) = 12.12, p = 0.001, η_p_^2^= 0.46), suggesting that posterior auditory regions might be especially sensitive to temporal structure in music. Posterior regions also responded more strongly overall to drums, as evidenced by a larger difference between intact melodic MIDI stimuli and intact drum stimuli in posterior compared to anterior regions (F(1, 14) = 28.97, p < 0.001, η_p_^2^ = 0.67). We also observed significant main effects of ROI (F(1,14) = 5.18, p = 0.04, η_p_^2^ = 0.27), hemisphere (F(1,14) = 11.30, p = 0.002, η_p_^2^ = 0.45) and a significant ROI x hemisphere interaction (F(1,14) = 12.80, p = 8.0e-04, η_p_^2^ = 0.48), but these are not relevant because they are due to overall differences in response magnitude across regions.

Together, these findings demonstrate that sensitivity to pitch and temporal pattern structure is strongly present in music-selective neural populations across hemispheres and different regions of auditory cortex, though posterior regions in both hemispheres may play a somewhat more prominent role in representing temporal musical structure.

#### Sensitivity to musical note patterns is concentrated in music-selective neural populations

Is sensitivity to note-pattern structure concentrated in music-selective neural populations or present in other nearby neural populations that also respond strongly to music? In our previous study, we found that voxel responses to a wide variety of natural sounds could be approximated by a set of six canonical response profiles, or “components”, one of which was music-selective. Four other components were found to reflect acoustic properties of the sound set (e.g., frequency, spectrotemporal modulation) and were concentrated in and around primary auditory cortex (PAC), consistent with prior results (Da Costa et al., 2011; Herdener et al., 2013; Hullett et al., 2016b; Humphries et al., 2010; Norman-Haignere & McDermott, 2018; Santoro et al., 2014b; Schönwiesner & Zatorre, 2009). Another component responded selectively to speech (Component 5) and clustered lateral to primary auditory cortex in the middle superior temporal gyrus.

We repeated our analyses for these other five canonical response components (**Figure 4A**). Some of these components showed substantial spatial overlap with music-selective neural populations and produced strong responses to many of the music stimuli tested. For example, Component 4, which shows sensitivity to pitch (Boebinger et al., 2021; Norman-Haignere et al., 2015), had weights that overlapped anterior music-selective neural populations and responded strongly to the melodic pitched MIDI music as well as speech and animal sounds, but only weakly to MIDI drums. Despite its strong response to music, we did not observe a significant effect of scrambling musical structure in the response of this pitch-sensitive component (repeated-measures ANOVA with factors “pitch-scrambling” and “time-scrambling” produced no significant main effects or interaction; all p’s ≥ 0.05, see **Table 1**). Thus, even though the spatial distribution of the pitch-sensitive and music-selective components substantially overlap, their response properties dissociate, highlighting the ability of voxel decomposition to isolate neural subpopulations within voxels that show different response characteristics.

**Figure 4.**
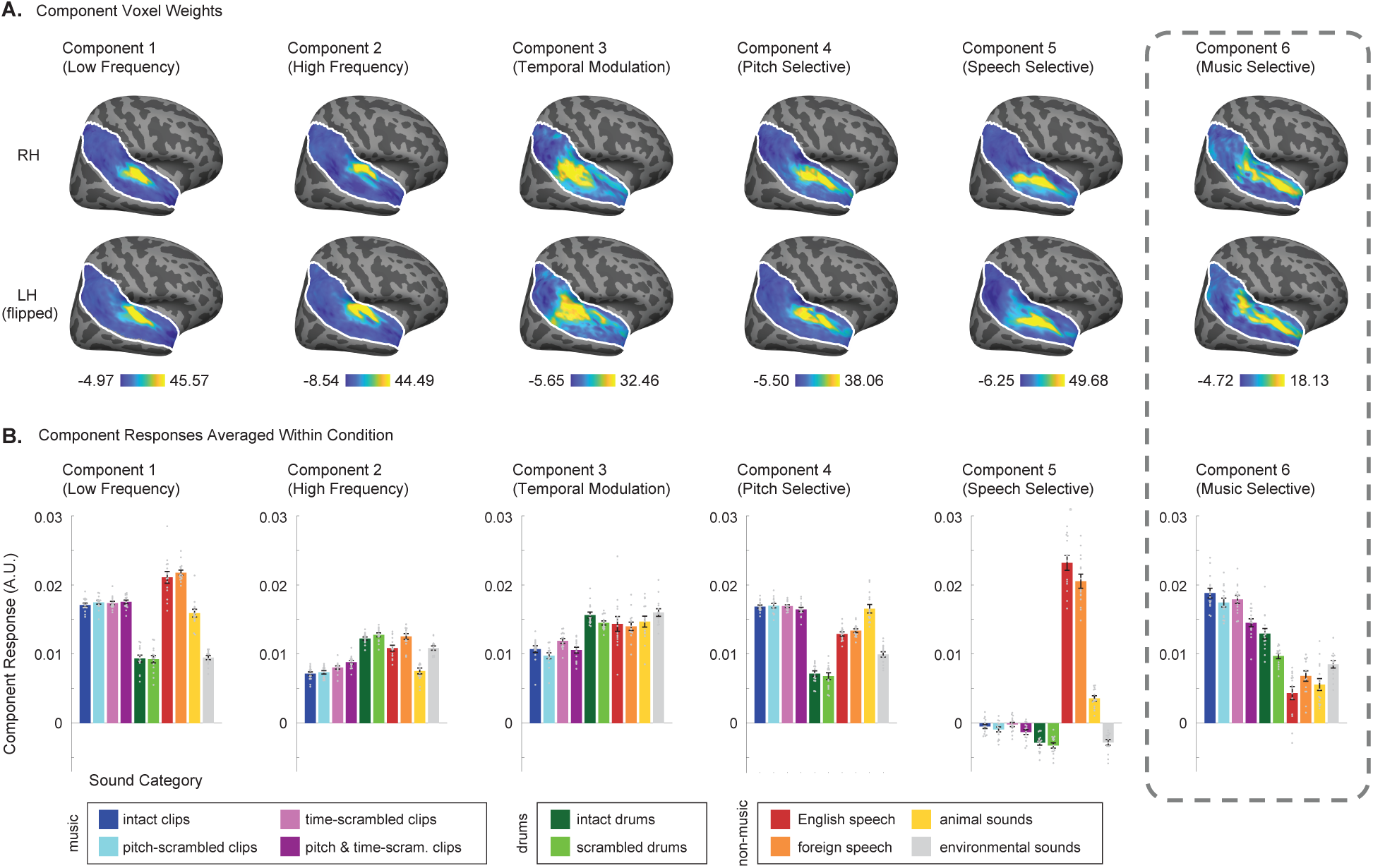
Response of all components to MIDI music and non-music control stimuli**. A.** Spatial distribution of component voxel weights for participants in the current study, inferred using the response components from Norman-Haignere et al. (2015). Voxel weights are averaged across participants in standardized anatomical coordinates. Color scale spans the central 95% of the weight distribution for each component. Music component voxel weights are the same as depicted in Figures 2B and **3A**. **B.** Component responses to the stimulus conditions in the main experiment, averaged across participants. Each gray dot corresponds to one participant’s component response to a given condition. Scatter plots are within-subject, such that differences between participants are removed by subtracting each participant’s mean response across conditions and then adding the global mean across participants. Error bars indicate ±1 within-subject SEM. Music component response is the same as depicted in Figure 2D.

We observed weak but significant effects of scrambling in some of the other remaining non-music components (see **Table 1**); however, the effects were much smaller than those in the music component in all cases (see **Table 2** for full ANOVA results; all 2-way and 3-way interactions involving “component” were significant: p’s < 0.01). We note that scrambling manipulations never perfectly match lower-level acoustic structure, and our component decomposition methods likely do not perfectly isolate music selectivity. It is thus not surprising that some non-music selective components show small effects of scrambling, and since component responses are highly reliable, small differences are able to reach statistical significance. Notably, the music-selective component was also the only component that showed evidence for a selective response to note-level structure. Specifically, the music component was the only component where the response to pitch and rhythm scrambled MIDI music was higher than the response to every non-music sound. Collectively, these results suggest that sensitivity to both note-level and note-pattern structure is concentrated in music-selective neural populations of anterior and posterior human non-primary auditory cortex, bilaterally.

## DISCUSSION

Our study demonstrates that non-primary neural populations just beyond primary auditory cortex show selectivity for the organization of musical notes across both pitch and time. These effects are concentrated in small, music-selective neural populations and are weak or absent in all other auditory populations, including nearby neural populations that overlap substantially with music-selective populations. Yet, within music-selective populations, selectivity for pitch and temporal pattern structure is broadly distributed across anterior and posterior regions and both hemispheres. Strong responses were also observed to scrambled music, which lacks coherent pitch and temporal patterns yet evoked substantially stronger activity than all other non-music sounds tested (speech, animal calls, environmental sounds). This suggests that music-selective populations are sensitive to both higher-order note patterns and to the acoustic properties that characterize individual notes (e.g., flat pitch contour, attack-sustain-decay temporal envelope). Together, these results suggest that representations of both single-note features and higher-order note patterns are concentrated in focal, bilateral music-selective neural populations.

### Selectivity for musical note and note-pattern structure in the auditory cortex

Prior work gives reason to think that auditory cortical representations may reflect sensitivity to the acoustic properties of individual musical notes. Non-primary neural populations selective for music and speech integrate information over 200-500 ms (Norman-Haignere, Long, et al., 2022), consistent with short-term representations of notes or syllables. Prior work has also found that flattening spoken pitch contours to be more note-like enhances responses in music-selective regions of auditory cortex (Sankaran et al., 2024; Tierney et al., 2013), supporting the idea that selectivity reflects tuning for structure on the level of individual notes. Under this hypothesis, musical pattern-level structure might be predominantly represented beyond the auditory cortex, for example, in frontoparietal regions that integrate across much longer timescales and show enhanced responses to surprising notes or chords (Levitin & Menon, 2003, 2005; Maess et al., 2001; Tillmann et al., 2003).

On the other hand, sensitivity to note patterns has been observed in the auditory cortex using a variety of methods, including decoding (Di Liberto et al., 2021; Hoefle et al., 2018; Lee et al., 2011; Schindler et al., 2013) and encoding (Di Liberto et al., 2020; Quiroga-Martinez et al., 2019; Sankaran et al., 2024) models, and inter-subject correlation (Farbood et al., 2015; Regev et al., 2021). Much of this prior work has not investigated whether this sensitivity to note patterns was specific to music or reflected more general acoustic processing. However, a recent intracranial study found that more surprising notes evoked larger responses in the auditory cortex, independent of generic acoustic or pitch features, and that this effect was correlated with an electrode’s overall selectivity for music vs. speech (Sankaran et al., 2024). Nevertheless, surprisal is just one aspect of musical structure, and it remains unclear how much of the cortical response to music is driven by surprisal compared to other forms of note pattern selectivity.

Our results suggest that both note-level and note-pattern structure contribute to auditory cortical representations of music, and that these representations are localized to small patches of music-selective neural populations. Music that preserved individual notes but disrupted tonal patterns elicited significantly lower responses than intact music, yet still evoked significantly stronger responses than all other non-music sounds tested. These findings are non-trivial in light of prior evidence that music-selective populations integrate information over relatively short timescales (on the order of a few hundred milliseconds; Norman-Haignere, Long, et al., 2022) and respond rapidly following the onset of musical structure (Norman-Haignere, Feather, et al., 2022). Although our results do not speak directly to the timescale of processing within music-selective neural populations, they demonstrate that their responses nonetheless depend not just on the presence of notes, but also on the patterned relationships among notes in both pitch and time.

While scrambled music differs acoustically from speech and other environmental sounds, prior work using more controlled manipulations suggests this difference cannot be explained by generic spectrotemporal tuning. For example, prior studies have found much stronger responses to natural music than spectrotemporally matched music in non-primary auditory cortex (Norman-Haignere & McDermott, 2018), and the magnitude of this difference is substantially larger than what we observed in the current study when comparing intact vs. scrambled music. This suggests that there is sensitivity to note-level structure that cannot be explained by generic acoustic features such as spectrotemporal modulation.

What types of musical pattern structure might underlie the selectivity we observe here? Although scrambling manipulations disrupt many musical features simultaneously and therefore cannot pinpoint which specific features drive the observed differences, they provide a powerful means of probing sensitivity to higher-order structure. Our manipulations preserved the acoustics of individual notes, while removing as much note-pattern structure as possible. Our pitch scrambling manipulation removed scale structure that characterizes the tonal structure of music, as well as statistics of melodic contour and voice-leading properties that characterize how typical tonal music unfolds over time. Our temporal scrambling manipulation removed both temporal periodicity and higher-order metrical structure. Importantly, both manipulations also affect consonance/dissonance, for example, by creating overlapping notes not present in intact music.

By defining broad constraints on the structure underlying music selectivity, our results lay the groundwork for future studies to test these representational hypotheses in greater detail using approaches such as encoding models.

### Methods for investigating sensitivity to musical structure

In this study, we used a scrambling manipulation inspired by prior work on visual and speech processing. The logic of this approach is that neural populations tuned to the statistical structure of natural stimuli should respond more weakly when that structure is disrupted (Liu et al., 2010; Mollica et al., 2020; Overath et al., 2015; Pallier et al., 2011). For example, the fusiform face area responds more strongly to intact than spatially rearranged faces (Liu et al., 2010), language regions in the frontal and temporal lobes respond more to sentences than to word lists (Mollica et al., 2020; Pallier et al., 2011), and regions of the superior temporal gyrus respond more to intact than scrambled speech (Overath et al., 2015). A few fMRI studies have applied scrambling techniques to music (Abrams et al., 2011; Farbood et al., 2015; Fedorenko et al., 2012; Norman-Haignere et al., 2015), but most of these studies scrambled natural audio recordings, which can introduce unnatural artifacts. By contrast, scrambling synthetic MIDI stimuli allowed us to largely preserve the acoustics of individual notes while selectively disrupting pitch and/or temporal pattern structure. Nonetheless, no scrambling method perfectly preserves all lower-level acoustic features, including those used here.

A second limitation of prior work is that music-selective neural populations were not explicitly isolated. For example, Fedorenko et al. (2012) found stronger responses to intact than scrambled music in scattered voxels across temporal and frontal lobes, but these voxels also responded robustly to visually presented sentences, suggesting they were not specifically tuned to music. Because music selectivity is weak in individual voxels, prior work suggests that component-based approaches are needed to isolate music-selective neural populations with fMRI. However, we note that our component methods are likely imperfect and do not completely isolate music selectivity in a single component. As a result, small scrambling effects occasionally appeared in non-music-selective components, but these effects were much weaker than in music-selective populations, even in components that spatially overlap substantially with music-selective neural populations. This pattern supports the conclusion that sensitivity to musical pattern structure is concentrated in focal music-selective neural populations rather than reflecting generic acoustic processing.

Alternative approaches for examining musical structure include violation paradigms and encoding models. Violation studies are often interpreted within predictive coding frameworks, in which neurons signal events that are unlikely given the prior context (e.g., out-of-key notes or chord) (Heilbron & Chait, 2017; Koelsch et al., 2019). These violations frequently activate frontal and parietal regions of the ventral attention network (Levitin & Menon, 2003, 2005; Maess et al., 2001; Tillmann et al., 2003), but these regions also respond strongly to non-musical violations (Doeller et al., 2003; Schubotz & von Cramon, 2002). Encoding models offer a complementary approach by quantifying the influence of specific features, such as surprisal, in natural music without highly unnatural violations (Di Liberto et al., 2020; Sankaran et al., 2024). To what extent can the results of our study be explained in terms of prediction error and musical surprisal? A simple unweighted prediction-error account would predict stronger responses to scrambled stimuli (Di Liberto et al., 2020; Koelsch et al., 2019; Sankaran et al., 2024; Vuust et al., 2022), contrary to our data. However, more nuanced predictive coding theories posit that neural responses reflect precision-weighted prediction errors, such that the influence of prediction errors depends on the reliability of the underlying predictive model (Barascud et al., 2016; Quiroga-Martinez et al., 2019; Sohoglu & Chait, 2016). Under this framework, extensive scrambling of musical structure may weaken the predictive model itself, reducing the precision assigned to prediction errors and thereby suppressing neural responses. This account predicts that local perturbations within otherwise coherent musical contexts would enhance responses, whereas global scrambling that disrupts the predictive structure of the stimulus would suppress them – a hypothesis that could be tested in future work.

Our scrambling methods provide one approach for separately investigating sensitivity to musical pitch (e.g., melody, harmony, tonality) and temporal structure (e.g., beat, rhythm), but are limited by the inherent entanglement of melody and rhythm. Drum stimuli partially address this issue, as they lack fine-grained note-level pitch, and thus help to isolate the contribution of temporal pattern processing in the absence of tonal structure. By contrast, isolating pitch pattern processing is more challenging because the constituent pitches of a melody necessarily unfold over time. Given this, our most direct measure of pitch pattern sensitivity is the contrast between time-scrambled and pitch-and-time-scrambled conditions, since neither contains coherent rhythm for pitch scrambling to disrupt. Overall, our results are consistent with the hypothesis that music-selective populations encode both temporal and pitch pattern structure, but we cannot firmly establish whether these dimensions are represented by overlapping or distinct neural populations, both because pitch and temporal structure are not cleanly separable in music and because fMRI provides a coarse measure of underlying neural activity.

The significant interaction between pitch and temporal scrambling (**Figure 2C**) indicates that the effect of scrambling in one domain was substantially reduced when the other domain remained intact, suggesting that the presence of any coherent note-pattern structure is sufficient to produce a comparatively strong response. Prior behavioral work shows that only a minimal amount of information is needed for listeners to instantiate complex tonal (Cuddy & Badertscher, 1987; Dowling, 1978; Oram et al., 1995; Smith & Schmuckler, 2004) and metrical structure (Brochard et al., 2003; Desain & Honing, 2003; Palmer & Krumhansl, 1990; Povel & Okkerman, 1981). This observation parallels findings in vision, where face-selective regions respond robustly whenever partial face structure is present (e.g. Arcurio et al., 2012; Tong et al., 2000).

### Functional organization of music selectivity in auditory cortex

Music-selective populations form at least two distinct, widely spaced clusters in anterior and posterior auditory cortex, and prior research suggests that there are prominent differences between these two regions. For example, anterior regions show more prominent tuning for fine spectral modulations characteristic of tonal sounds, while more posterior regions show tuning for rapid temporal modulations characteristic of impulsive sounds (e.g., drums) (Hamilton et al., 2018; Hullett et al., 2016b; Norman-Haignere et al., 2015; Santoro et al., 2014b; Schönwiesner & Zatorre, 2009). Anterior auditory cortex also contains neural populations that respond to tones with resolved harmonics that have a strong sense of pitch and show sensitivity to pitch changes (Norman-Haignere et al., 2013; Patterson et al., 2002). Although our scrambling manipulation preserved spectrotemporal modulation, it disrupted pitch and temporal patterning, making it plausible that anterior regions would favor pitch structure and posterior regions would favor temporal structure. Our results instead show sensitivity to both pitch and temporal structure in both clusters, though posterior music-selective regions responded more strongly overall to drum stimuli, consistent with sensitivity to rapid temporal modulations characteristic of impulsive sounds (Hamilton et al., 2018; Hullett et al., 2016b; Norman-Haignere et al., 2015; Santoro et al., 2014b; Schönwiesner & Zatorre, 2009). They also showed a stronger time-scrambling effect than anterior music-selective populations, consistent with a small posterior bias for temporal pattern sensitivity. Thus, while our results rule out the possibility of a strong anatomical anterior-posterior division between temporal and pitch pattern selectivity, they leave open the possibility that there might be weak biases aligned with this distinction.

Prior studies have also proposed that there are hemispheric asymmetries, such that the right hemisphere is specialized for pitch/melody and the left hemisphere for rhythm (Albouy et al., 2020; Zatorre & Belin, 2001). This account predicts stronger pitch-scrambling effects in the right hemisphere and stronger time-scrambling effects in the left hemisphere. Contrary to this prediction, we observed no hemispheric differences in the effects of either type of scrambling. Instead, sensitivity to both pitch and temporal pattern structure was evident in both hemispheres, consistent with a broadly distributed representation of musical structure. At the same time, this sensitivity remains highly focal, confined to small clusters of music selectivity that also track note-level structure.

Behavioral ratings of musicality correlated with the response profile of the music component (**Figure S1**), suggesting these populations are related to the perception of music. At the same time, music perception likely involves other brain regions, including those beyond the auditory cortex that may not be specific to music per se. For example, the perception of metrical structure may be influenced by motor regions (Brett & Grahn, 2007; Fedorenko et al., 2012; Janata et al., 2002; Lee et al., 2011; Matthews et al., 2020; Williams et al., 2021), frontal regions that code expectation may modulate activity in the auditory cortex (Koelsch et al., 2019), and limbic regions likely play an important role in linking musical representations with memory and emotional structures.

### Limitations and future directions

Like all studies, our conclusions are limited by the resolution of the methods used to measure neural responses. Functional MRI pools activity from hundreds of thousands of neurons and thus inevitably reflects mixed selectivities, even using component methods. Methods with higher spatial and temporal resolution have been able to infer a larger number of reliable response components that reflect finer-grained patterns of selectivity across sounds (Norman-Haignere, Feather, et al., 2022). High temporal resolution might thus be required to observe dissociations between the processing of pitch and temporal structure in music, as suggested by another intracranial EEG study that found differences in the temporal dynamics of expectations associated with note pitches vs. onset times (Di Liberto et al., 2020), which would not be detectable with fMRI. As the precision of human neural recordings continues to advance (Leonard et al., 2024), it may become possible to identify finer subdivisions within music-selective cortex, deepening our understanding of the neural computations that support human music perception.

## ACKNOWLEDGEMENTS

This study was supported by the National Science Foundation (BCS-1634050 to J.H.M. and BCS-2342908 to S.N.H.) and the National Institutes of Health (NIDCD-R00-DC018051 to S.N.H., NIDICD-R01DC017970 to J.H.M., NICHD-DP1HD091947 to N.G.K., and OD-S10OD021569).

## AUTHOR CONTRIBUTIONS

D.L.B.: Conceptualization, Data Curation, Formal Analysis, Investigation, Methodology, Project Administration, Validation, Visualization, Writing – original draft, Writing – review & editing

J.H.M: Conceptualization, Funding Acquisition, Methodology, Resources, Software, Supervision, Writing – review & editing

N.G.K.: Conceptualization, Funding Acquisition, Resources, Software, Supervision, Writing – review & editing

S.V.N.-H: Conceptualization, Funding Acquisition, Methodology, Supervision, Writing – review & editing

