## Supplemental Figure 1 for "Music-selective cortex is sensitive to structure in both pitch and time"

**SUPPLEMENTARY MATERIAL**

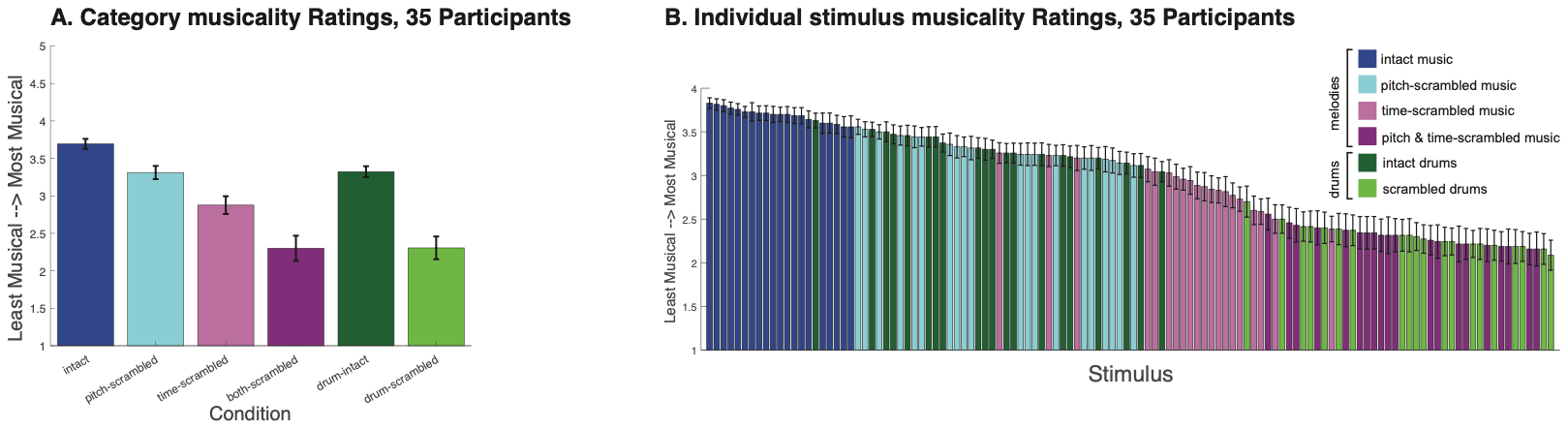

**Figure S1.** An online experiment was run on Amazon’s Mechanical Turk in which participants (n = 35) listened to each of the 80 MIDI melodies and 40 MIDI drum stimuli and rated them on a scale of 1 (“not musical at all”) to 5 (“very musical”). **A.** Musicality ratings averaged over categories. **B.** Musicality ratings for each individual stimulus. Error bars indicate ±1 within-subject SEM.
