## Supplemental Table 1 for "Music-selective cortex is sensitive to structure in both pitch and time"

**SUPPLEMENTARY MATERIAL**

| Localizer | Version 1  *(Norman-Haignere et al., 2015)* | Version 2  *(Boebinger et al., 2021)* | Version 3  *New subjects* |
| --- | --- | --- | --- |
| MRI scanner | Trio | Prisma | Trio |
| Head coil | 32-channel | 32-channel | 32-channel |
| TR | 3.4 | 3.4 | 3.59 |
| TA | 1 | 1.02 | 1.19 |
| TE | 30ms | 33ms | 30ms |
| Flip angle | 90 | 90 | 90 |
| Number of slices | 15 (partial coverage) | 48 (whole-brain) | 45 (whole-brain) |
| Slice thickness | 4 mm | 3 mm | 2.8 mm |
| Slice gap | 10% | 10% | 10% |
| Matrix | 96 x 96 | 96 x 96 | 96 x 96 |
| Voxel size | 2.1 x 2.1 x 4 mm | 2.1 x 2.1 x 3 mm | 2 x 2 x 2.8 mm |
| Simultaneous multi-slice acceleration (SMS) | none | 4 | 3 |
| Number of stimuli | 165 | 192 | 30 |
| Number of stimulus repetitions in each “mini-block” | 5 | 3 | 3 |
| Number of stimulus blocks per run | 15 | 24 | 30 |
| Run duration | 5.4 min | 5.5 min | 6.5 min |
| Number of runs | 33 | 48 | 10 |
| Number of stimulus repetitions | 3 | 6 | 10 |
| Number of participants | 2 | 7 | 6 |

**Table S1.** Scan parameters for the three versions of the component localizer scan sessions.
